# Changes in the Physicochemical Properties and Microbial Communities of Rhizospheric Soil after Cassava/Peanut Intercropping

**DOI:** 10.1101/570937

**Authors:** Xiumei Tang, Saiyun Luo, Zhipeng Huang, Haining Wu, Jin Wang, Guoying Shi, Liangqiong He, Faqian Xiong, Jing Jiang, Jing Liu, Guojian Liao, Ronghua Tang, Longfei He

## Abstract

Cassava/peanut intercropping is a popular cultivation method in southern China and has the advantages of apparently increased yield and economic efficiency compared with monoculture, however, the ecological benefits of this method are poorly understood. This study aimed to investigate the effects of intercropping on the physicochemical properties and microbial community structures of soil. Field trials were performed to determine the effects of cassava/peanut intercropping on rhizospheric soil nutrient content, enzyme activities, microbial quantity and microbial community structure. The microbial community was characterized by 16S rRNA tag-based high-throughput sequencing on the Illumina MiSeq platform. Results showed that cassava/peanut intercropping could improve the physicochemical properties of rhizospheric soil by increasing the available nutrient content, pH, bacterial quantity, and some enzyme activities and by altering the microbial community structure. 16S rRNA gene sequencing demonstrated that the microbial community structure varied between the intercropping and monoculture systems. Nitrospirae, Verrucomicrobia and Gemmatimonadetes were more abundant in the intercropping system than in the monocultures. Redundancy analysis (RDA) revealed that the abundances of *DA101*, *Pilimelia* and *Ramlibacter* were positively correlated with environmental parameters such as available nitrogen and pH, and these were dominant genera in the rhizospheric soil of the intercropped peanut plants.

## INTRODUCTION

Intercropping is a global agricultural practice involving the growing of two or more crops at the same yield during the growing season, and this method has become increasingly important for the improvement of soil quality and to increase crop productivity [**1-3**]. Intercropping has additional advantages, such as efficient acquisition of nutrients, establishment of soil microbial diversity and improved utilization of land resources [**4, 5**]. There have been several intercropping systems developed to date, such as maize/peanut, wheat/cotton, and wheat/maize [**6-8**].

It has been reported in previous studies that intercropping can change the soil microecology, as indicated by increasing farmland biodiversity [**9**]. Intercropping can effectively improve the mobilization and uptake of nitrogen (N), phosphorus (P), potassium (K), and micronutrients via interspecific interactions in the rhizosphere[**4, 10, 11**]. In addition, legume/cereal intercropping systems could improve the utilization of phosphorus (P) by root exudation of organic acids from legume crops also improve legume nitrogen (N) uptake by enhanced nodulation of legume crops [**12, 13**].

Soil microbe and soil enzyme activities play important roles in nutrient cycling, organic matter decomposition and suppression of soil-borne pathogens in the rhizosphere ecosystem [**14-16**]. Plants can release root exudates, thereby affecting the rhizosphere microbial community [**17**]. The potential activities of soil enzymes can be influenced by changes in microbial community composition. Due to the quantitative and qualitative differences between the root exudates of intercropping and monocropping systems, differences in the microbial communities can be observed [**18**]. Many studies have investigated the changes in the biochemical and microbial characteristics of rhizospheric soils caused by intercropping [**1, 19**]. For the alfalfa/rye intercropping system, it has been found that intercropping can affect the soil microbial composition and soil enzymatic activities [**20**]. By using phospholipid fatty acid (PLFA) analysis, it was found that the soil urease and invertase activities and the soil gram-negative (G-) bacterial abundance increased significantly in the peanut/*Atractylodes lancea* system [**21**]. Many studies have shown that the maize/peanut intercropping system can facilitate the acquisition of Fe and Zn by the peanut crop and can improve the yield of both crops [**10, 22**].

Cassava/peanut intercropping is a typical intercropping cultivation mode in southern China because the original spacing of the cassava crop remains unchanged when intercropped with peanut, thereby providing a distinct yield advantage. However, at present, although much research has been conducted regarding the selection of the cassava/peanut intercropping model and the associated yield benefits, the corresponding theoretical research remains insufficient. Previous studies on cassava/peanut intercropping have mainly focused on the uptake and utilization of nutrients, photosynthesis, agronomic traits, yield, efficiency and nutrient conversion efficiency in the soil [**23, 24**]. However, the influence of the soil microecological environment in the cassava/peanut intercropping system remains unknown. The objectives of this study were to determine the effect of the cassava/peanut interaction patterns on physiochemical properties and microbial community, and to examine the relationships among the microbial communities, soil enzymatic activities, and soil nutrients.

## MATERIALS AND METHODS

### Materials

The tested cassava crop, named “Huanan 205”, was provided by farmers in Wuming County of Guangxi Province and is a main cassava variety in Guangxi, China. The tested high-yield and shade-tolerant peanut crop, named “Guihua 836”, was provided by the Cash Crops Research Institute of the Guangxi Academy of Agricultural Sciences and is suitable for intercropping.

### Experimental site and soil

The field trial was conducted in the Lijian Scientific Base of the Guangxi Academy of Agricultural Sciences, Nanning City, Guangxi province in China (23°14′N, 108°03′E), at an altitude of 99 m above sea level. The field site had been previously used for monoculturing cassava.The tested soil was acid red loam, the organic matter content, total nitrogen content, total phosphorus content, total potassium content, available nitrogen content, available phosphorus content and available potassium content of which were 16.2 g/kg, 1.34 g/kg, 0.53 g/kg, 12.6 g/kg, 70.5 mg/kg, 13.9 mg/kg and 98 mg/kg, respectively. The pH value was 5.8.

### Experimental design and management

On March 8, 2016, cassava and peanut were planted simultaneously in the field. Monocultured cassava(MC) and monocultured peanut(MP) crops were the controls, and cassava/peanut intercropping was the treatment group, which were intercropped cassava(IC) and intercropped peanut(IP). For monoculture, cassava was planted with a row spacing of 1.1 m × 0.8 m and with equivalent line spacing. For monoculture, peanut was planted in narrow-wide row spacing. The line spacing for peanut in the wide line was 0.5 m. The row spacing of the peanut plants in the narrow row was 0.3 m × 0.16 m. For cassava/peanut intercropping, two lines of peanut were planted between two lines of cassava. The line spacing between cassava and peanut was 0.4 m. The row spacing for cassava and peanut intercropping was 1.1 m × 0.8 m and 0.3 m × 0.16 m, respectively. The experiment was arranged in plots (6 m×8 m) in a randomized design with three replicates in each treatment.

Cassava and peanut respectively received different fertilizing amount in seed furrow according to their growth demand before sowing. All peanut treatments received 450 kg. ha^−1^ compound NPK granulated fertilizers (N-P_2_O_5_-K_2_O=15-15-15) and 750 kg. ha^−1^ fused calcium-magnesium phosphate fertilizer ∈ available P_2_O_5_ 18%), and all cassava treatment only received 750 kg. ha^−1^ compound NPK granulated fertilizers. The crops were irrigated two times during crop growth, based on crop water requirement and soil water content. Cultivating and weeding were done about two month after sowing, and made sure that the experimental plot have not weed and prevent soil compaction.

### Soil sampling

On July 10, 2016 (when mature peanut plants were ready for harvest), ten plants of cassava and peanut per sample were uprooted. The rhizospheric soil from loose soil and cohesive soil from the plant roots (loose soils were shaken off and cohesive soils were brushed with a sanitized soft brush)were collected, mixed and separated into three sealed virus-free bags, which were kept in an icebox and taken to the laboratory. One bag was maintained in the refrigerator at 4°C and used for soil microbe determination. One bag was maintained in the refrigerator at −80°C and used for extracting soil DNA and for high-throughput sequencing. The last bag was dried naturally, ground and sieved for determining the nutrient content and soil enzyme activity.

### Soil chemical analysis

The nutrient content was measured according to the Technical Specifications for Soil Analysis. The available N, available P, available K and organic matter levels were measured by the alkaline hydrolysis diffusion method, sodium bicarbonate extraction/Mo-Sb colorimetry, ammonium acetate extraction/flame photometry and the potassium bichromate titrimetric method, respectively.

Soil enzyme activity was measured by colorimetry and titration [**25**]. Catalase activity, sucrase activity, proteinase activity, urease activity and acid phosphatase activity were measured by permanganate titration, sodium thiosulfate titration, ninhydrin colorimetry, indophenol blue colorimetry and the disodium phosphate benzene colorimetric method, respectively.

### Determination of soil microbial quantity

Soil microbial quantity was measured by the conventional microculture method [**26**]. Bacteria, fungi and actinomycetes were cultured in beef extract-peptone medium, Martin medium and Gao 1 medium, respectively. The Shannon-Wiener index method was used to calculate the biodiversity index (H): H = −Σ(ni/N)×ln(ni/N) [**27**].In this formula, ni is the microbial quantity of species i, and N is the total microbial quantity.

### Soil DNA extraction, PCR amplification, high-throughput sequencing and analysis

Total genomic DNA was extracted from the samples using the FastDNA SPIN Kit (MP Biomedicals, Santa Ana, USA) according to the manufacturer’s instructions. DNA concentration and purity were monitored on 1% agarose gels, and the DNA was diluted to 1 ng/μL using sterile water. The DNA extracts were stored at −80°C until subsequent PCR amplification. The total genomic DNA was subjected to PCR amplification using the 515f/806r primer pair, which amplifies the V4 region of the 16S rDNA gene [**28**], following a previously described protocol [**29**]. All PCRs were carried out in 30-μL reactions with 15 μL of Phusion^®^ high-fidelity PCR master mix, 0.2 μM forward and reverse primers, and approximately 10 ng of template DNA. Thermal cycling consisted of an initial denaturation at 98 °C for 1 min; which was followed by 30 cycles of denaturation at 98 °C for 10 s, annealing at 50 °C for 30 s, and elongation at 72 °C for 30 s; and a final step at 72 °C for 5 min. The PCR products were mixed with equal volumes of 1× loading buffer (containing SYBR Green), and electrophoresis was conducted on a 2% agarose gel for detection. Samples with a bright band at 400-450 bp were chosen for further experiments. The PCR products were mixed at equal concentrations. Then, the mixture of PCR products was purified with the GeneJET Gel Extraction Kit (Thermo Scientific). Sequencing libraries were generated using the NEB Next^®^ Ultra− DNA Library Prep Kit for Illumina (NEB, USA) following the manufacturer’s recommendations, and the index codes were added. The library quality was assessed on a Qubit 2.0 fluorometer (Thermo Scientific) and an Agilent 2100 bioanalyzer system. Finally, the library was sequenced on an Illumina MiSeq platform by the Novogene Corporation (Beijing, China), and 250/300-bp paired-end reads were generated.

Paired-reads from the original DNA fragments were merged based on a previously described method [**30**]. Sequencing reads were assigned to each sample according to the individual unique barcodes. Sequences were analyzed with the QIIME (Quantitative Insights Into Microbial Ecology) software package and UPARSE pipeline [**31**]. The reads were first filtered by QIIME quality filters. Default settings for Illumina processing in QIIME were used. Then, the UPARSE pipeline was used to select operational taxonomic units (OTUs) at 97% similarity. For each OTU, a representative sequence was selected and used to assign taxonomic composition by using the RDP classifier [**32**]. Then, the estimated species richness was determined by rarefaction analysis [**33**]. Redundancy analysis (RDA) was performed to analyze the correlation between environmental factors and microbial communities.

### Data analysis

The means, standard deviations and histograms of the four treatments were plotted using GraphPad Prism (version 5.0). One-way variance analysis was performed with SPSS18.0, and Duncan’s methods were used to test the homogeneity of variance at confidence levels of 0.01 and 0.05.

## RESULTS

### Effects of cassava/peanut intercropping on the soil nutrient content

Fig. **1** a-h shows that the total N, total K, available N, available K, and organic matter content and the pH value in the rhizospheric soil of the intercropped peanut plants were greater than those of the monocultured peanut plants. The total N and available N content and the pH value for the intercropped peanut plants were significantly different from those for the monocultured peanut plants, increasing by 18.1%, 9.1% and 19.6%, respectively, whereas the total P and available P levels for the intercropped peanut plants were lower than those for the monocultured peanut plants.

**Fig 1.**
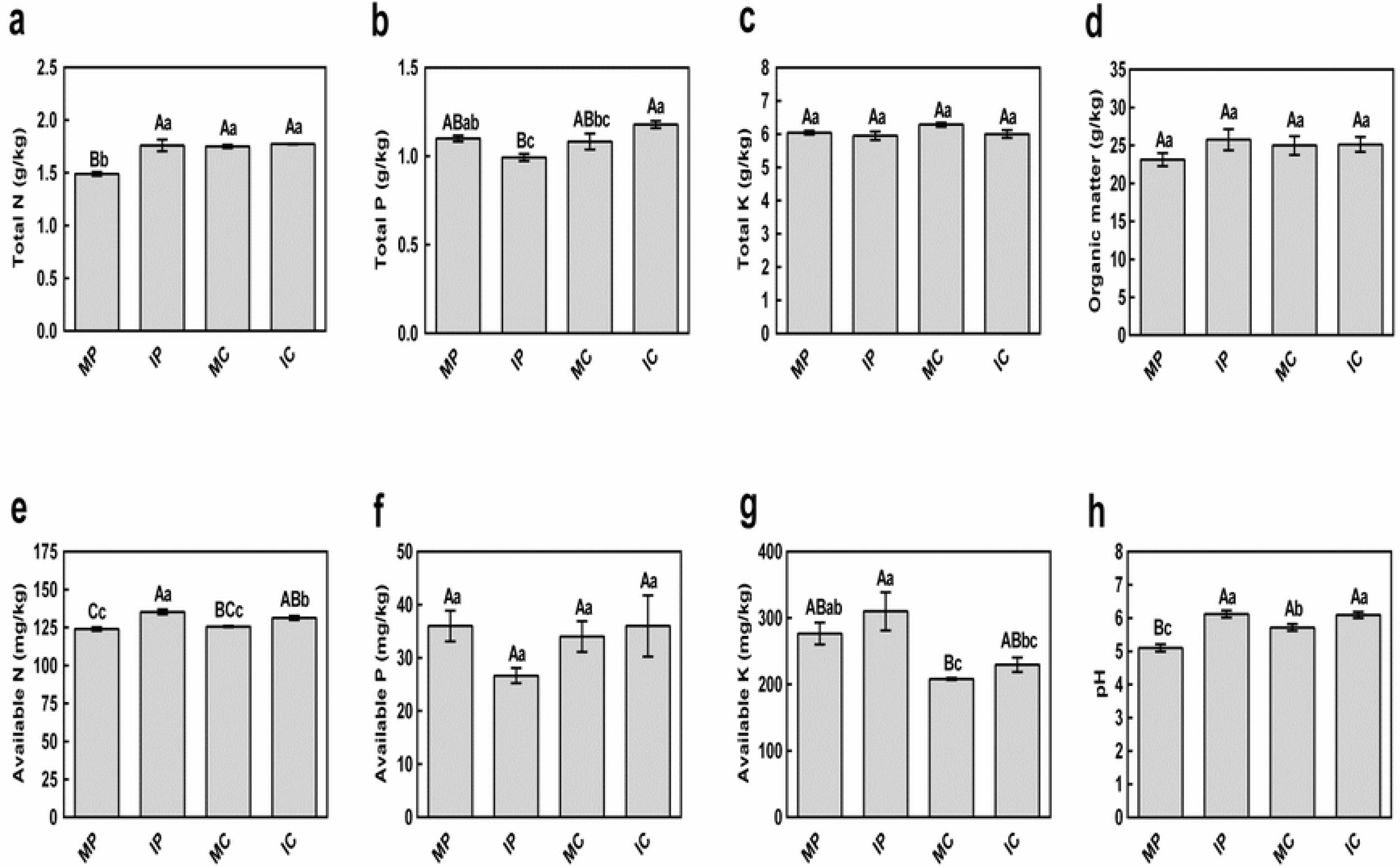
Effect of the cassava/peanut intercropping system on the physical and chemical properties of the soil. a: percentage of total nitrogen in the rhizospheric soil; b: percentage of total phosphorus in the rhizospheric soil; c: percentage of total potassium in the rhizospheric soil; d: organic matter content in the rhizospheric soil; e: available nitrogen content in the rhizospheric soil; f: available phosphorus content in the rhizospheric soil; g: available potassium content in the rhizospheric soil; h: pH of the rhizospheric soil environment. MP: peanut in monoculture; IP: peanut intercropped with cassava; MC: cassava in monoculture; IC: cassava intercropped with peanut. Bars with different capital letter and small letters within a substrate are significantly different at P<0.01 and P<0.05, respectively.

All the nutrient indices in the rhizospheric soil of the intercropped cassava plants (total N, total P, total K, available N, available P, available K and organic matter content and pH value) were greater than those in the rhizospheric soil of the monocultured cassava plants. The total P and available N content and the pH value for the intercropped cassava plants were significantly different from those for the monocultured cassava plants, increasing by 9.3%, 4.5% and 7.0%, indicating that cassava/peanut intercropping could improve the soil pH, increase the nutrient content and markedly boost the total N and available N content in the peanut rhizosphere as well as the total P and available N content in the cassava rhizosphere. Intercropping greatly affected the available N content in the peanut and cassava rhizospheres. However, compared to the levels in the monocultured peanut and cassava rhizospheres, the available P content in the rhizospheric soils of intercropped cassava and peanut plants exhibited contrasting trends.

### Effects of cassava/peanut intercropping on the soil enzyme activity

The catalase, sucrase, urease and acid phosphatase activities in the rhizospheric soil of the intercropped peanut plants were all higher than those in the rhizospheric soil of the monocultured peanut plants, increasing by 2.8%, 10.1%, 24.2% and 24.7%, respectively, while the proteinase activity in the rhizospheric soil of the intercropped peanut plants did not differ significantly from that in the rhizospheric soil of the monocultured peanut plants, decreasing by 10.4% (Fig. **2**).

**Fig 2.**
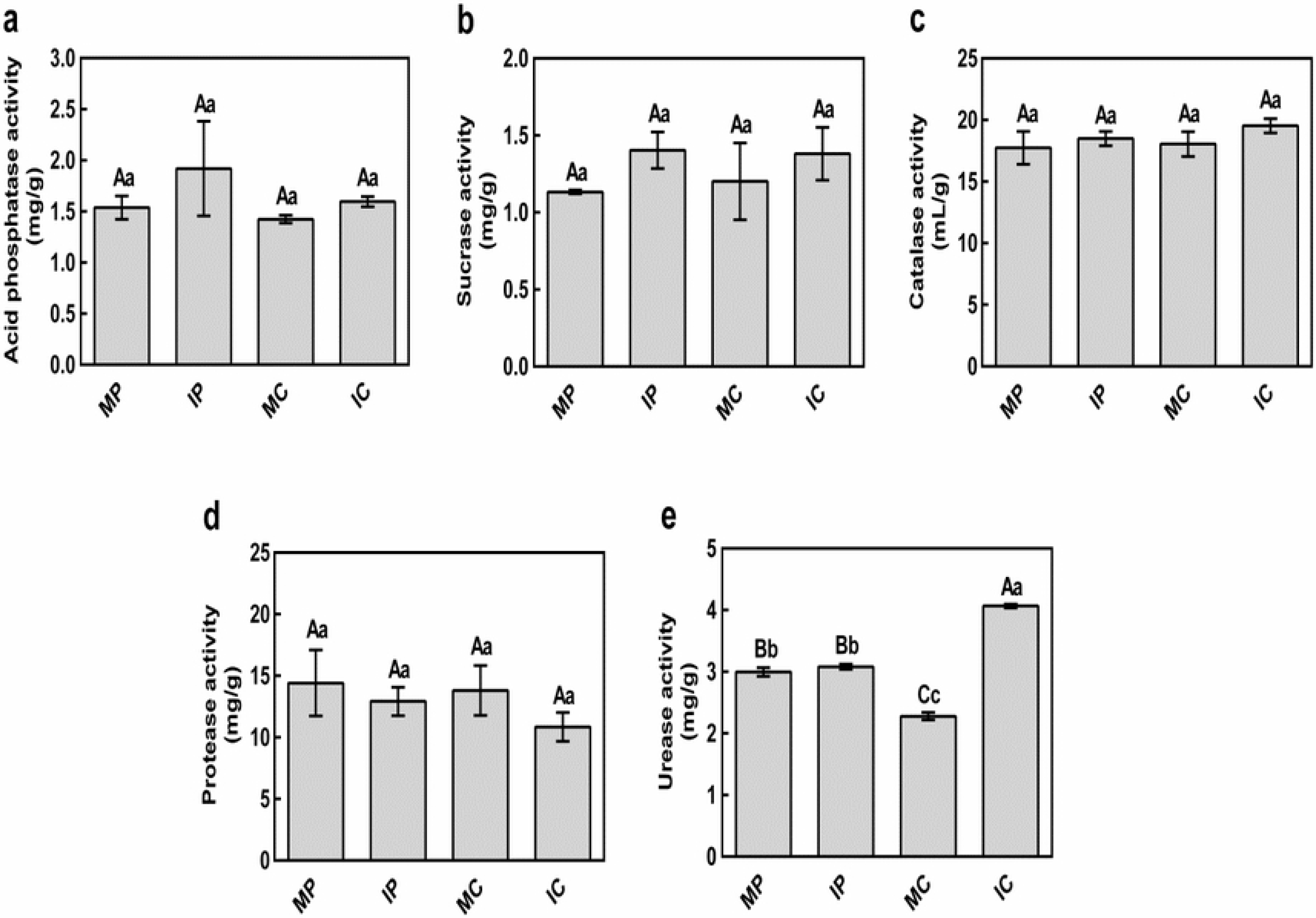
Effect of cassava/peanut intercropping on soil enzyme activities. MP: peanut in monoculture; IP: peanut intercropped with cassava. MC: cassava in monoculture; IC: cassava intercropped with peanut. Bars with different capital letter and small letters within a substrate are significantly different at P<0.01 and P<0.05, respectively.

Similar to the results for the intercropped peanut plants, the catalase, sucrase, urease and acid phosphatase activities in the rhizospheric soil of the intercropped cassava plants were all higher than those observed for the monocultured cassava plants, of which the difference in urease activity between intercropped cassava (78.5% increase) and monocultured cassava was significant. The proteinase activity in the rhizospheric soil of intercropped cassava did not differ significantly from that observed for monocultured cassava, decreasing by 21.5%(Fig. **2**).

Compared to monoculture, cassava/peanut intercropping improved the catalase, sucrase, urease and acid phosphatase activities; however, intercropping reduced the proteinase activity. Among the abovementioned enzyme activities, urease activity was the one that was most affected in the rhizospheric soil.

### Effects of cassava/peanut intercropping on the microbial quantity in the rhizospheric soil

The bacterial abundance (27.5% increase), fungal abundance (2.5% increase), actinomycete levels (53.1% increase), total microbial quantity (31.7% increase) and microbial diversity index (9.3% increase) in the rhizospheric soil of the intercropped peanut plants were all greater than those observed for the monocultured peanut plants. Among these parameters, the bacterial abundance and actinomycete levels increased greatly, and the difference was significant (Fig. **3a-e**).

**Fig 3.**
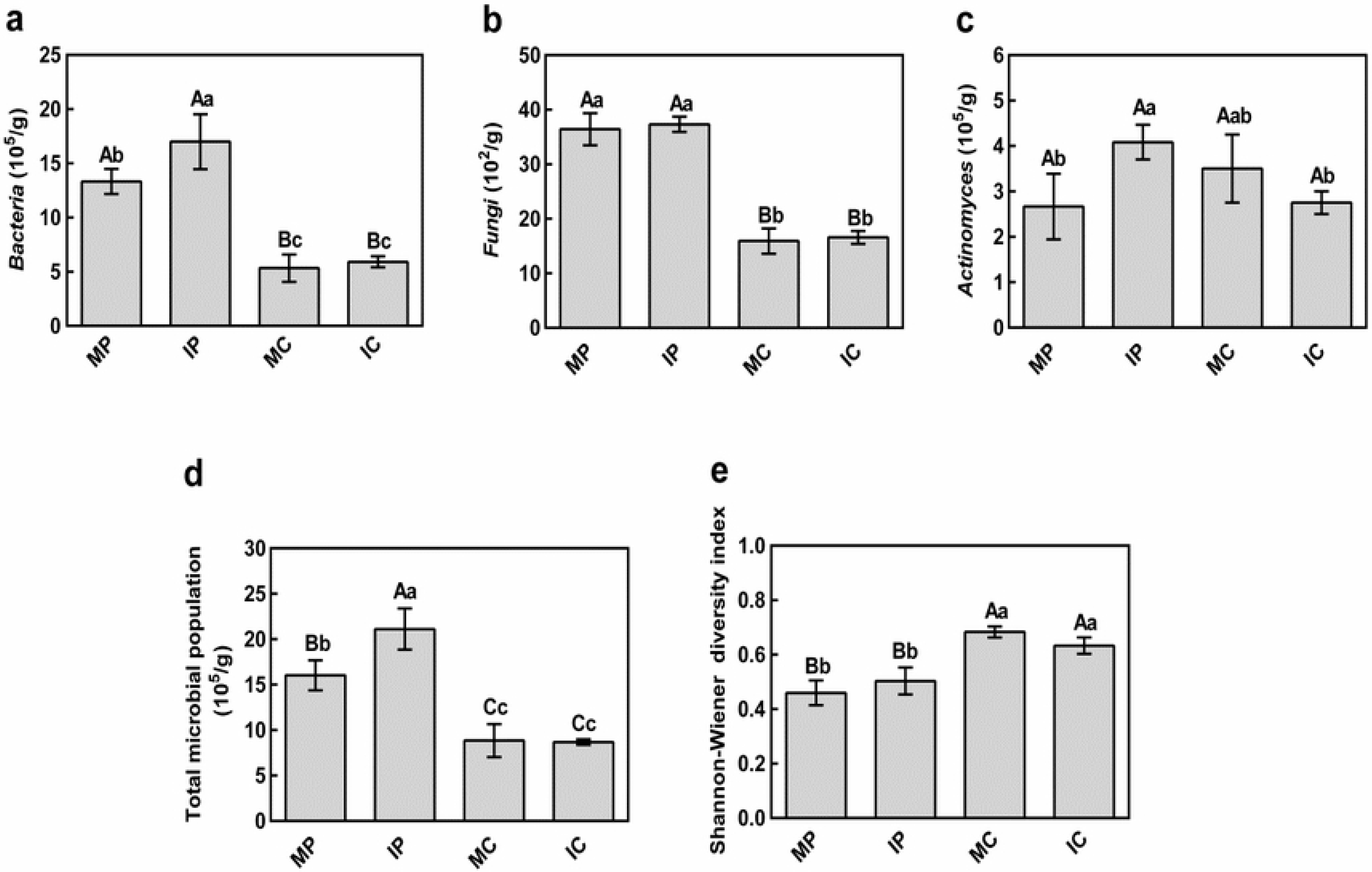
Effect of cassava/peanut intercropping on soil microbial quantity. MP: peanut in monoculture; IP: peanut intercropped with cassava; MC: cassava in monoculture; IC: cassava intercropped with peanut. Bars with different capital letter and small letters within a substrate are significantly different at P<0.01 and P<0.05, respectively.

The bacterial abundance (10.9% increase) and fungal abundance (2.5% increase) in the rhizospheric soil of intercropped cassava were slightly greater than those observed for monocultured cassava, while the actinomycete levels, total microbial quantity and microbial diversity index were marginally lower in the rhizospheric soil of intercropped cassava than those observed for monocultured cassava. The total microbial quantity and microbial diversity index did not differ significantly between intercropped cassava and monocultured cassava. It can be deduced that cassava/peanut intercropping is much better than monoculture in terms of increasing the bacterial abundance and actinomycete levels in the peanut rhizospheric soil. However, the cassava/peanut intercropping system barely affects the total microbial quantity in the cassava rhizospheric soil, demonstrating that the total microbe amounts in the rhizospheric soils of intercrops vary with crops (Fig. **3**).

### Effects of cassava/peanut intercropping on the microbial community

16S rRNA gene sequencing on the Illumina MiSeq platform yielded a total of 43,519 valid sequences, which represented the wide diversity of the microbial community. Sequence analyses at the phylum and genus taxonomic levels are shown in Fig. **4** and Fig. **5**, respectively. The taxonomic distribution at the phylum level is described in Fig. **4**, Proteobacteria was the most abundant phylum in all the samples, accounting for 28.24 to 37.45% of the total valid reads in all the samples, with an average relative abundance of 34.45%. Actinobacteria was the second most abundant phylum in all the samples, with an average relative abundance of 20.70%. The other dominant phyla were Acidobacteria (12.55–18.32%, with an average value of 14.79%), Chloroflexi (7.19–8.40%, with an average value of 7.76%), Gemmatimonadetes (3.80–5.83%, with an average value of 4.63%), Nitrospirae (3.27–5.81%, with an average value of 4.16%), Planctomycetes (1.60–4.36%, with an average value of 2.96%), Verrucomicrobia (1.46–4.51%, with an average value of 2.71%), and Bacteroidetes (1.03-3.75%, with an average value of 1.91%). In addition, Nitrospirae, Verrucomicrobia and Gemmatimonadetes were more abundant the rhizospheric soils of the peanut and cassava plants in the intercropping system than in the soil of the monocropping systems. Bacteroidetes and Planctomycetes were also more abundant in the rhizospheric soil of the peanut plants of the intercropping system than in those of the monocropping system, but these phyla were less abundant in the rhizospheric soil of the monocultured cassava plants than in that of the intercropped cassava plants. Other phyla, such as Proteobacteria, Actinobacteria, Acidobacteria and Chloroflexi did not exhibit significantly different abundances between the monoculture and intercropping systems.

**Fig 4.**
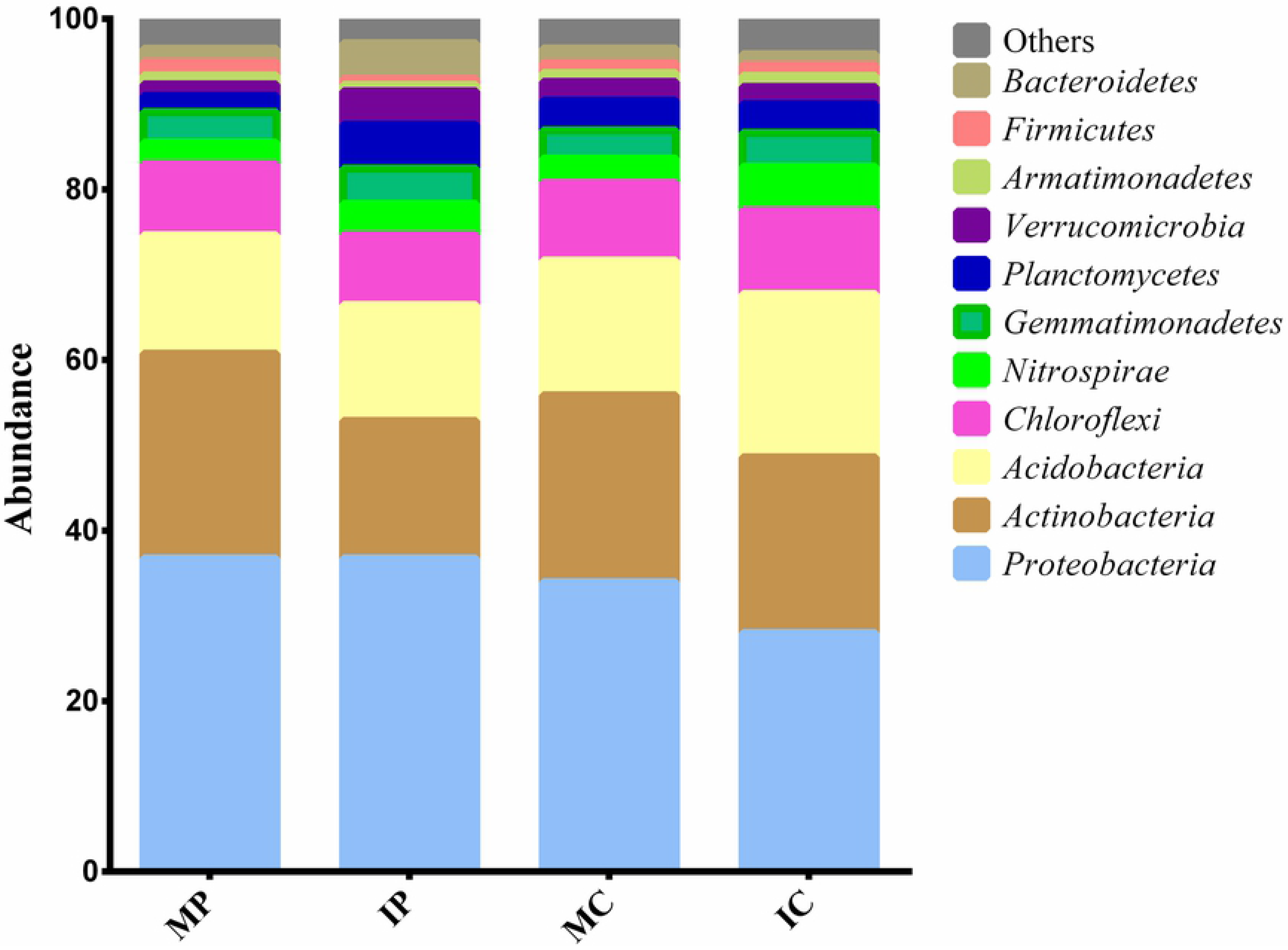
Taxonomic classification of bacterial reads from different soil environments under different planting patterns at the phylum level using RDP classifier. MP: peanut in monoculture; IP: peanut intercropped with cassava; MC: cassava in monoculture; IC: cassava intercropped with peanut.

**Fig 5.**
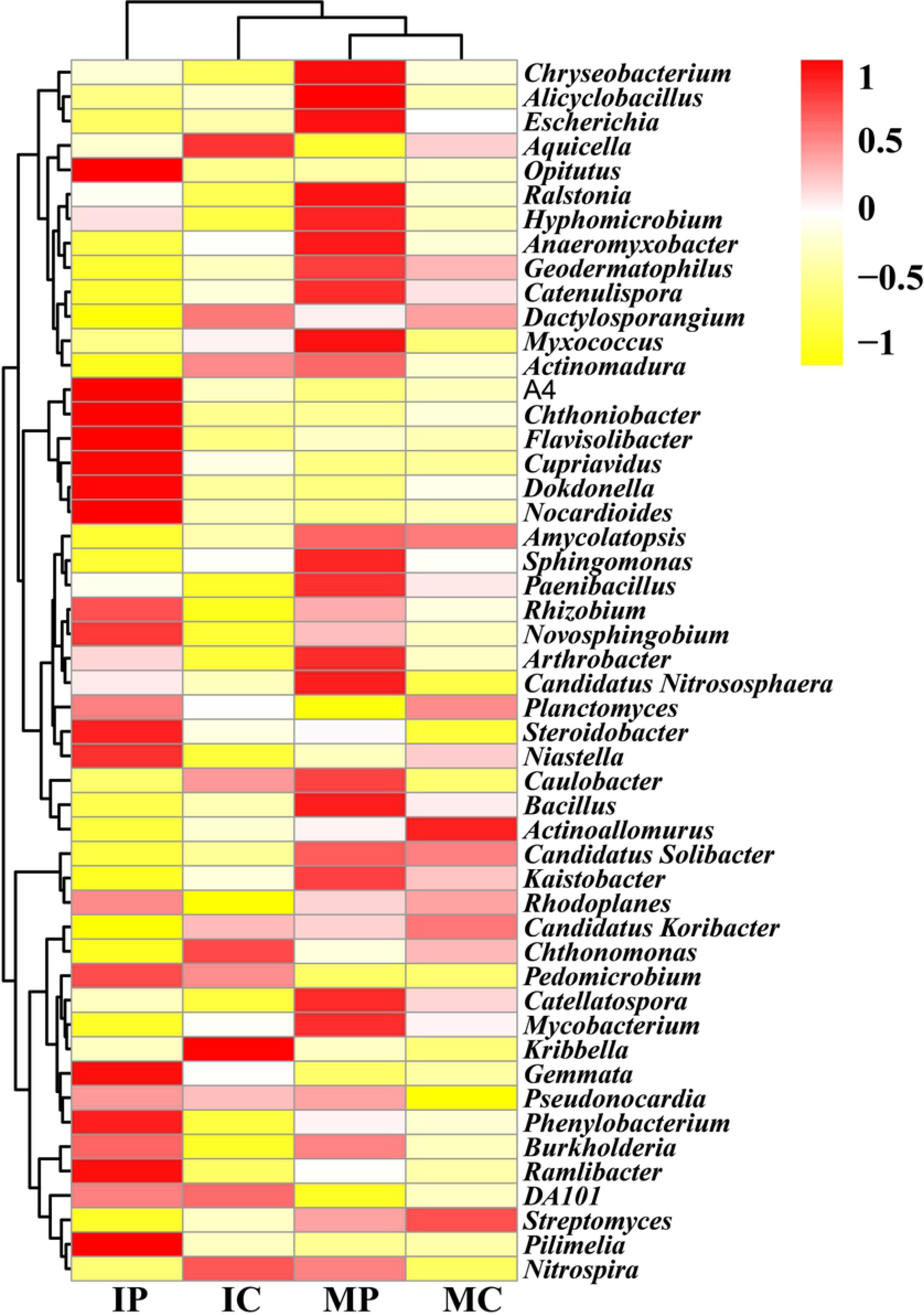
Heatmap analysis of the distribution of dominant genera among the four treatments. The double hierarchical dendrogram shows the microbial distribution among the four treatments. The relative values for the microbial genera are indicated by color intensity; the legend can be found at the top of the figure. The abundance is expressed as the value of the targeted sequence to the total number of high-quality sequences from each soil sample. MP: peanut in monoculture; IP: peanut intercropped with cassava; MC: cassava in monoculture; IC: cassava intercropped with peanut.

The most abundant genera within the four samples in different soil environments and under different planting patterns were also determined, as depicted in Fig. **5**. The different soil samples exhibited different dominant genera, for instance, *Aquicella, Chthonomonas, Kribbella, DA101* and *Nitrospira* were more abundant in the rhizospheric soil of the cassava plants in the intercropping system, but *Actinoallomurus* and *Streptomyces* were more abundant in the cassava monoculture system. Compared with the cassava rhizospheric soil, the peanut rhizospheric soil exhibited a high diversity of dominant genera. There were 15 dominant genera in the rhizospheric soil of the intercropped peanut plants, including *Optitutus*, *A4*, *Chthoniobacter*, *Flavisolibater*, *Dokdonella*, and *Pilimelia*, but these genera were not highly abundant in the peanut monoculture system, which had different dominant genera, such as *Chryseobacterium, Alicyclobacillus, Escherichia, Ralstonia,* and *Hypomicrobium*. Therefore, it was clearly demonstrated that cassava/peanut intercropping could change the microbial community structures, and the microbial communities of the intercropping system were distinct from those of the monocropping system. Furthermore, the microbial communities in the rhizospheric soils of the same crop can differ due to different planting patterns such as monoculture and intercropping.

Principal component analyses showed that samples from monoculture and intercropping systems were separated from each other, even though the monocultured peanut and intercropped peanut were distributed in quadrant 2 and quadrant 1, 4 respectively(fig. **6**), which were the same crop in different planting patterns, suggesting that different planting patterns favor different microbial communities. This finding may be associated with the environmental parameters in the different planting patterns. Therefore, correlations between environmental factors and the core genera were determined by RDA.

**Fig 6.**
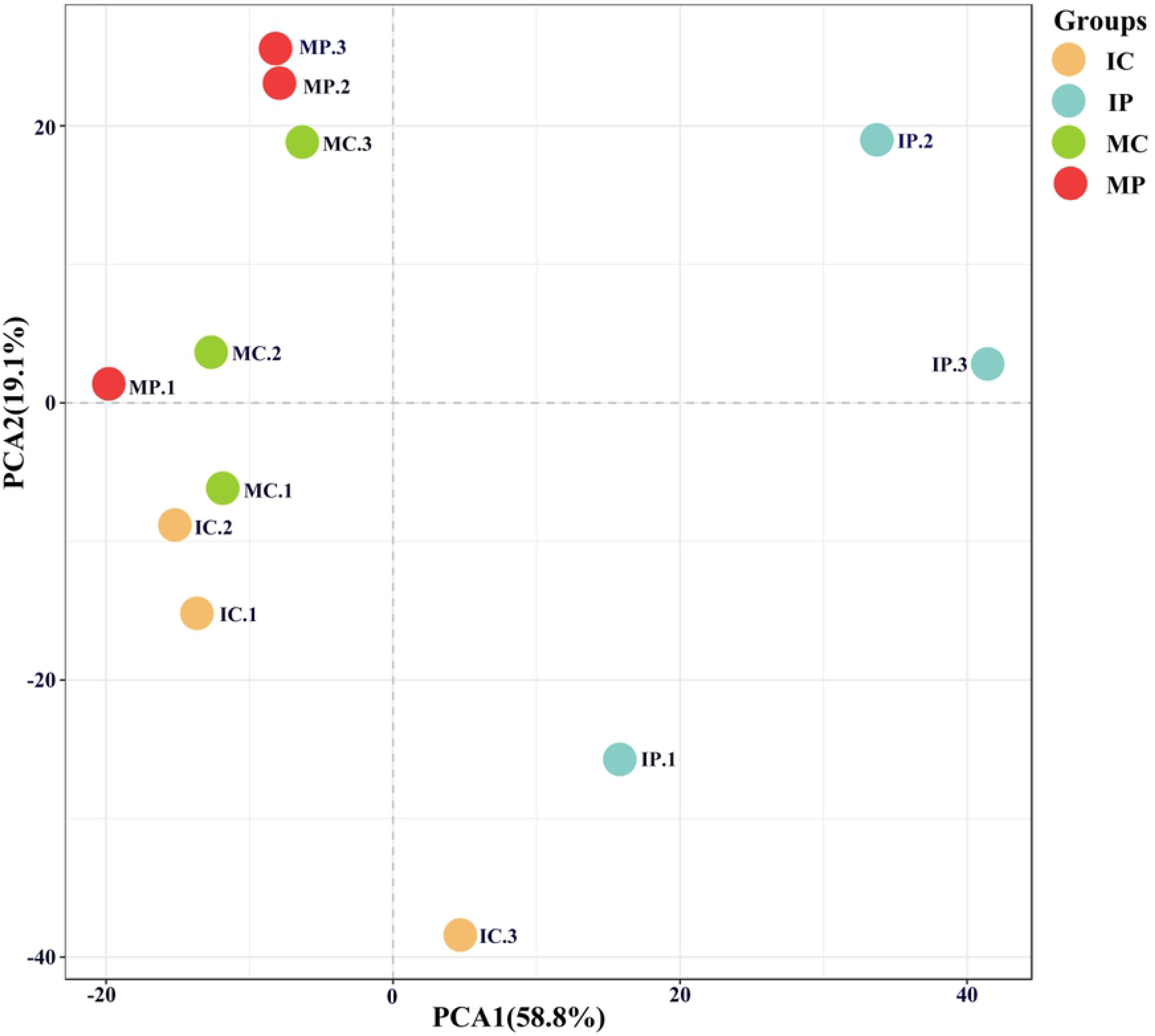
Principal component analyses of bacterial communities in the monoculture and intercropping systems based on Euclidean distance at the OTU level. MP: peanut in monoculture; IP: peanut intercropped with cassava; MC: cassava in monoculture; IC: cassava intercropped with peanut.

### Redundancy analysis (RDA) of microbial communities and environmental parameters

Microbial communities exhibit high correlation with intrinsic environmental parameters. Therefore, RDA was used to examine the mechanism via which microbes can adapt to the changes in physicochemical environments in situ. Correlations between the important environmental parameters and the microbial community was discerned by RDA, as shown in Fig. **6**. The length of the arrow corresponding to an environmental parameter indicates the strength of the environmental parameter in relation to the overall microbial community.

The RDA suggested that there were differences in the bacterial communities in the different planting patterns. The results demonstrated that available nitrogen, catalase, organic matter, sucrase activity, acid phosphatase activity, urease activity, total nitrogen, pH, total potassium and available potassium were positively correlated with the RDA axis 1 and were strongly and significantly associated with the overall microbial community. In contrast, the total phosphorus, available phosphorus and protease activity were negatively correlated with the RDA axis 1. The results revealed that available nitrogen, pH, catalase activity and sucrase activity had the greatest impact on the microbial community. Additionally, the abundances of some microbial genera, such as *DA101*, *Pilimelia*, and *Ramlibacter* (Fig. **7**), were positively correlated with available nitrogen. These were also the dominant genera among intercropped peanut plants (Fig. **4**).

**Fig 7.**
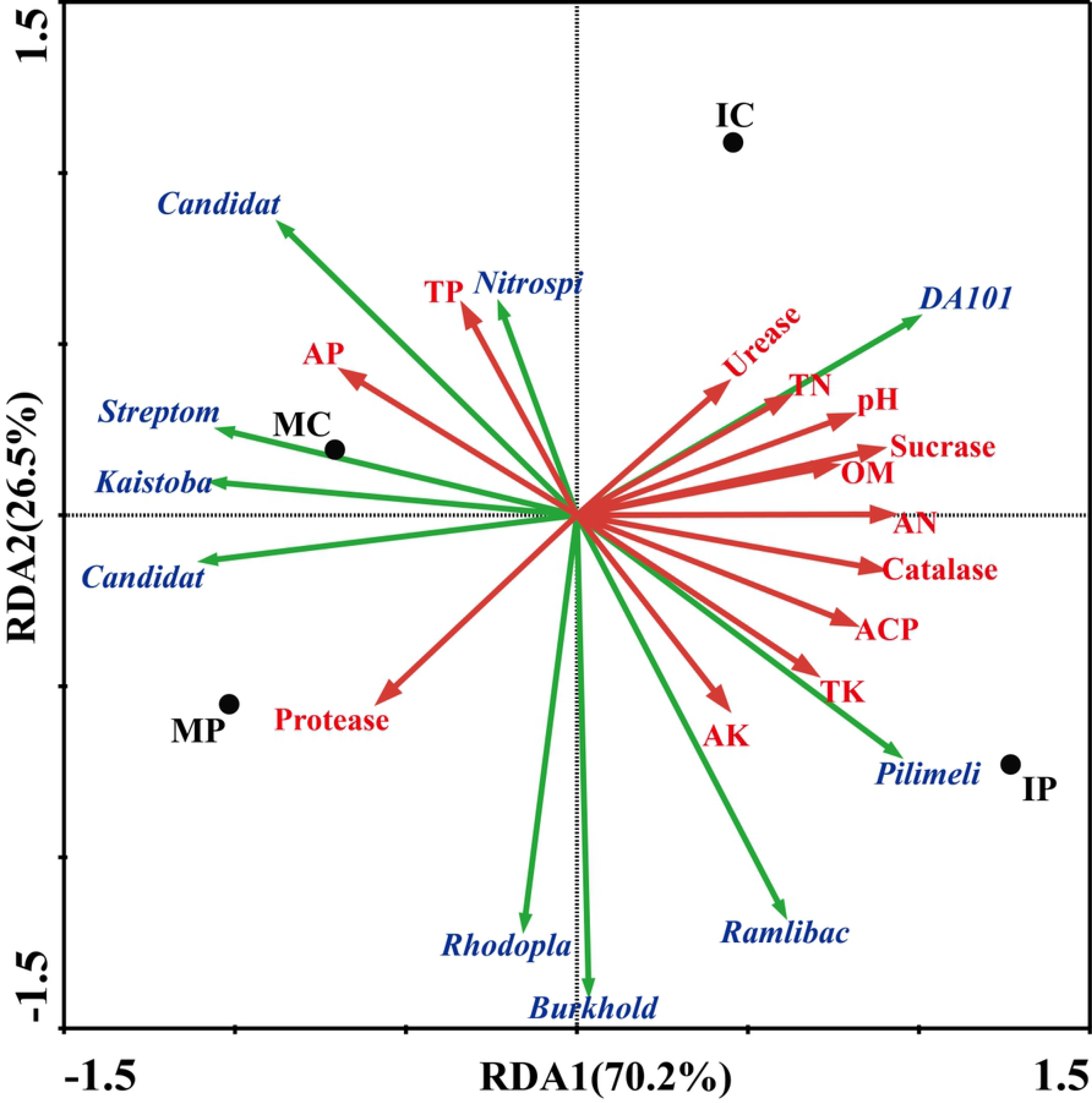
Redundancy analysis (RDA) of 16S rRNA gene data and environmental parameters. Arrows indicate the direction and magnitude of the environmental parameters associated with bacterial community structure. MP: peanut in monoculture; IP: peanut intercropped with cassava; MC: cassava in monoculture; IC: cassava intercropped with peanut.

## DISCUSSION

### Variations in soil nutrient content in the cassava/peanut intercropping system

Soil nutrients are essential for crop growth. The results of this study prove that cassava/peanut intercropping can improve the soil in terms of pH and nutrient levels. In particular, this intercropping system can incredibly increase the available N content. The available P levels in intercropped peanut and cassava exhibited different variation trends, and the available P content was higher in intercropped cassava than in monocultured cassava; however, the available P content was lower in intercropped peanut than in monocultured peanut. This finding was probably associated with competition for P uptake between peanut and cassava in the intercropping system. Previous studies on sugarcane/soybean [**34**], chestnut/tea [**35**], maize/peanut [**36**], soybean/tea [**37**] and maize/pepper [**38**] intercropping have also shown that intercropping can boost the soil nutrient content. Based on the results of previous research reports and the current study, there are three possible factors that can account for the increase in soil nutrient content in the cassava/peanut intercropping system: root exudates, microbial function and enzymatic activity, and biological nitrogen fixation by intercropped peanut plants. Root exudates can improve the effective nutrient content because they can change physicochemical conditions of the rhizospheric soil and activate soil nutrients by dissolving nutrients that are difficult to dissolve [**39**]. The soil nutrient content can also increase due to increased microbial diversity together with increased catalase and acid phosphatase activity, as shown in Fig. **2a**, **2c** and Fig. **3d**, **3e**, which can improve the efficiency of the transformation of organic nutrients to effective inorganic nutrients. Moreover, microbial community maybe play an important role in improving soil nutrient of intercropping system. The phylum Nitrospirae, Verrucomicrobia and Gemmatimonadetes were more abundant in the intercropping soil than monocropping systems(Fig. **4**). The genera *DA101*, *Pilimelia* and *Ramlibacter* were positively correlated with available nitrogen (Fig.**7**). Therefore, the mechanisms of interaction between microbial community and soil nutrient should be well worthy to be studied further.

### Variations in soil enzyme activity in the cassava/peanut intercropping system

Soil enzymes, which are involved in the cycling of nutrients and materials, are associated with soil nutrients and microbes [**40, 41**]. The results showed that cassava/peanut intercropping improved the activities of catalase, sucrase, urease and acid phosphatase but reduced the proteinase activity. Among all the abovementioned enzymatic activities, urease activity was the most affected in the cassava rhizospheric soil. With regard to soil enzymatic activity in the intercropping system, the results from previous research reports have not been consistent. Kuang Shizi et al. found that compared to monocultured banana, intercropped banana (banana/soybean, banana/peanut, banana/ginger) could increase the activities of urease, sucrase and acid phosphatase, but the catalase activity decreased [**42**]. This result was not consistent with the result of the present study, which showed that cassava/peanut intercropping could improve catalase activity. Chai Qiang et al. [**43**] reported that compared to monocultured maize, the activities of urease and acid phosphatase decreased markedly in maize/chick pea intercropping. In contrast, Fruit/grass (grain) intercropping enhanced the activities of urease, acid phosphatase and catalase [**44**]. Based on the abovementioned studies, it can be concluded that soil enzymatic activity varies with enzyme type, crop cultivar and intercropping mode. Further research should be conducted on the variation mechanisms of soil enzyme activities and on the interactions of these enzymes with microbes and nutrients in soil.

### Variations in soil microbial quantity in the cassava/peanut intercropping system

Soil microbes, which are important for the formation of agricultural ecosystems and determine soil fertility and quality, play a key role in crop residue degradation, humus formation, and nutrient conversion and cycling [**45**]. Compared to monocultured peanut, the total microbial quantity in the rhizospheric soil was higher for intercropped peanut, for which the amounts of bacteria and actinomycetes were greatly increased. This result was consistent with the results obtained for pear/aromatic plants, sugarcane/soybean, wheat/cotton and maize/soybean intercropping systems [**34, 36, 46, 47**]. Lu Yahai and Zhang Fusuo [**48**] stated that root exudates and crop residues could create a beneficial environment and provide carbon and energy sources for soil microbes. It can be inferred that intercropping leads to richer root exudates and crop residues than monoculture because there is more than one crop in the same plot in an intercropping system. An increased supply of energy sources for rhizospheric microbes leads to an improved growth environment, increasing the total microbial quantity. Additionally, the microbial diversity index of intercropped peanut was greater than that of monocultured peanut, while the microbial diversity index of intercropped cassava was marginally lower than that of monocultured cassava. Different reports have made different statements regarding variations in soil microbial diversity. Zhang [**9**] found that microbial diversity in the rhizospheric soil of crops increased with the increasing of aboveground crop cultivars. In contrast, Dong Yan and Song Yana showed that rhizospheric microbial diversity in a wheat/bean intercropping system was lower than that for monocultured bean at the mid-late stage [**49, 50**], which was inconsistent with the results of this study, which found that cassava/peanut intercropping could boost microbial diversity in the rhizospheric soils of both intercropped and monocultured peanut. Therefore, soil microbial diversity is likely affected by crop combinations, crop varieties, growth period, root exudates, soil environment and sampling time.

### Changes in the microbial community in the cassava/peanut intercropping system

Microbes are important indicators of changes in soil ecosystems, and the soil microbial quantities and community structures are influenced by the environment, tillage and planting patterns [**7**, **51, 52**]. In the present study, Proteobacteria, Actinobacteria, Acidobacteria, Gemmatimonadetes, Nitrospirae, Planctomycetes, Verrucomicrobia, and Bacteroidetes were the dominant phyla in the cassava/peanut intercropping system. These phyla have been shown to be common phyla in farmland soils [**53, 54**]. The most dominant bacteria in the intercropping system were Proteobacteria, which is consistent with the results of a summary report for several soils [**55**]. The members of Proteobacteria exhibit wide morphological and metabolic diversity, playing crucial roles in the global carbon, nitrogen and sulfur cycles [**56, 57**]. In addition, Nitrospirae, Verrucomicrobia and Gemmatimonadetes were more abundant in the intercropping system than in the monocropping systems, and these phyla are also well represented in agricultural ecosystems [**58**]. Members of the bacterial phylum Verrucomicrobia have been detected in agricultural grassland soils [**59**] and black soils [**60**] and play important roles in biogeochemical cycling processes; these bacteria are strongly influenced by the soil pH and C:N ratio [**61**]. Members of the phylum Gemmatimonadetes have been found in various soils and have the ability to accumulate polyphosphates [**62, 63**].

The phylum Nitrospirae consists of gram-negative bacteria that act as nitrifiers, oxidizing nitrites to nitrates. Consequently, the increased abundance of the three dominant phyla might be associated with carbon, nitrogen and phosphorus nutrient cycling and soil acidity, which affects the nitrogen and phosphorus content in the intercropping system. Our results showed that intercropping could simultaneously increase the abundances of the three dominant phyla (Fig. **4**) and pH, available nitrogen, and available phosphorus content of the cassava rhizospheric soil (Fig. **1**). The mechanism for this effect remains unclear and should be explored in future research. However, several bacterial phyla varied between the intercropped peanut and intercropped cassava compares with the monocultured systems. For instance, Bacteroidetes and Planctomycetes were more abundant in intercropped peanut than in monocultured peanut but were less abundant in monocultured cassava than in intercropped cassava.

Similarly, a heatmap analysis of the dominant genera also showed that different soil samples exhibtied different dominant genera between the intercropping and monoculture systems. *Aquicella, Chthonomonas, Kribbella, DA101* and *Nitrospira* were more abundant in the rhizospheric soil of intercropped cassava, while the rhizospheric soil of intercropped peanut exhibited a number of different dominant genera compared with intercropped cassava, including *Optitutus*, *A4*, *Chthoniobacter*, *Flavisolibater*, *Dokdonella*, and *Pilimelia*, suggesting that differences in planting patterns and crop rhizospheres can explain differences in microbial community structures. This result might be associated with the effect of rhizosphere interactions and root exudates of different crops in intercropping systems on the physicochemical properties of the soil.

### Correlations between environmental parameters and the microbial community

Soil microbial community structures are always associated with the physicochemical properties of the soil. He et al. demonstrated that soil properties (such as pH and moisture content) significantly influence microbial species richness, composition and structure, which may determine or modify the functioning of the ecosystem [64, 65]. By RDA of the relationships between microbial communities and environmental parameters, Xu et al. also indicated that, based on the DGGE profile, community structures are highly correlated with the concentrations of pollutants (deca-BDE and octa-BDE) and soil properties (total nitrogen content) [66, 67].

By RDA, we identified additional important soil factors that affect bacterial community structures. The results indicated that urease activity, catalase activity, sucrase activity, total nitrogen, pH, organic matter, available nitrogen, acid phosphatase activity, total potassium and available potassium were positively correlated with the RDA axis 1 and were strongly and significantly correlated the overall microbial community. In contrast, total phosphorus, available phosphorus and protease activity were negatively correlated with the RDA axis 1. The results also revealed that available nitrogen, pH, catalase activity and sucrase activity had the greatest impact on the microbial community. The results are similar with those of previous studies. For example, by studying the effect of applied urea on rhizospheric soil bacterial communities in a greenhouse assay, Shang and Yi [**57**] reported that the bacterial communities were significantly correlated with the available nitrogen content and pH of the soil. Additionally, the abundance of *DA101*, *Pilimelia*, and *Ramlibacter* were positively correlated with important environment parameters such as available nitrogen and pH. These related genera were also dominant in the intercropped peanut plants, which may explain the increase in available nitrogen in the intercropping system. The abundance of the genus *DA101*, belonging to the phylum Verrucomicrobia, increased in the soil of the intercropping system, which was reported to be significantly negatively correlated with total nitrogen content and pH [**61**]. These results indicate that high N content is favorable for *DA101* growth, which is similar to our results, in which *DA101* abundance was seen to be significantly positively correlated with available nitrogen content, which is consistent with changes in the abundance of the phylum Verrucomicrobia in the intercropping system (Fig. **4**). Recently, Brewer et al. [**68**] assembled a draft genome of an organism named *Candidatus* Udaeobacter copiosus, which is a representative of the *DA101* clade, and speculated that *Ca*. U. copiosus is an oligotrophic soil bacterium that reduces its requirement for soil organic carbon by acquiring costly amino acids and vitamins from the environment. *Pilimelia* and *Ramlibacter* are seldom reported in agricultural soil, and their functions of these bacteria remain unclear; therefore, further research was performed to determine the correlations of these two genera with available nitrogen and other soil factors.

## CONCLUSIONS

(1) Cassava/peanut intercropping could increase the available N content, pH, and total N content of peanut rhizospheric soil as well as the pH, total P content and available N content of cassava rhizospheric soil. No significant difference in K content was observed between the cassava/peanut intercropping system and the monocultures.
(2) Compared to monoculture, cassava/peanut intercropping increased the activities of catalase, sucrase, urease and acid phosphatase but decreased the proteinase activity. Urease activity was most affected in the rhizospheric soil of intercropped cassava and was significantly different between intercropped cassava and monocultured cassava.
(3) In the intercropping system, microbial quantity and biodiversity index increases in the peanut rhizosphere soil; in particular, the abundances of bacteria and actinomycetes were much greater in the rhizospheric soil of intercropped peanut than in that of monocultured peanut. In the cassava rhizosphere soil, the abundances of bacteria and fungi for intercropped cassava were slightly greater than those for monocultured cassava, while the actinomycete levels, total microbial quantity and microbial diversity index were marginally lower for intercropped cassava than for monocultured cassava. None of the microbial factors differed significantly between intercropped cassava and monocultured cassava.
(4) The most dominant phylum in the cassava/peanut intercropping system was Proteobacteria, and Nitrospirae, Verrucomicrobia and Gemmatimonadetes were more abundant in the intercropping system than in the monocultures. At the genus level, the dominant genera varied among the different planting pattern and crop species. Notably, the abundances of *DA101*, *Pilimelia*, *Ramlibacter* were positively correlated with the microbial communities and environmental parameters such as available nitrogen and pH, suggesting that the three genera played an active role in shaping the intrinsic microbial communities and changing the nutrient nitrogen cycle. However, the mechanisms associated with the interactions between soil physicochemical factors and microbial communities need to be studied in further detail.

## ACKNOWLEDGMENTS

We would like to thank the National Natural Science Foundation Project (31401318 & 31660371), Technology System for Modern Agriculture (CARS-13-Southern China), Guangxi Natural Science Foundation Project (2017GXNSFAA198144) and Scientific Project from the Guangxi Academy of Agricultural Sciences (2015JZ12 &2017JZ12) for financial support.

## REFERENCES

1. Zhou X, Yu G, and Wu F. Effects of intercropping cucumber with onion or garlic on soil enzyme activities, microbial communities and cucumber yield. Eur J Soil Biol. 2011; 47: 279–287.

2. Zhang F, Li L. Using competitive and facilitative interactions in intercropping systems enhances crop productivity and nutrient-use efficiency. Plant Soil. 2003; 248: 305–312.

3. Vandermeer JH. The Ecology of intercropping. Cambridge: Cambridge University Press; 1992. pp. 237.

4. Li L, Tilman D, Lambers H, Zhang FS. Plant diversity and overyielding: insights from belowground facilitation of intercropping in agriculture. New Phytol. 2014; 203: 63–69.

5. Lacombe S, Bradley RL, Hamel C, Beaulieu C. Do tree-based intercropping systems increase the diversity and stability of soil microbial communities. Agr Ecosyst Environ. 2009; 131: 25–31.

6. Awal MA, Koshi H, Ikeda T. Radiation interception and use by maize/peanut intercrop canopy. Agr Forest Meteorol. 2006; 139: 74–83.

7. Zhang L, vander WW, Zhang S, Li B. Effects of intercropping on the quality of cotton were minor and mostly below detection threshold. Field Crop Res. 2007; 103 (3): 178–188.

8. Yang CH, Chai Q, Huang GB. Root distribution and yield responses of wheat/maize intercropping to alternate irrigation in the arid areas of northwest China. Plant soil environ. 2010; 56 (6): 253–262.

9. Zhang FS, Shen JB, Li L, Liu X. An overview of rhizosphere processes related with plant nutrition in major cropping systems in China. Plant Soil. 2004; 260: 89–99.

10. Inal A, A. Gunes F, Zhang Z, Cakmak I. Peanut/maize intercropping induced changes in rhizosphere and nutrient concentrations in shoots. Plant Physiol Biochem. 2007; 45: 350–356.

11. Zuo YM, Zhang FS. Effect of peanut mixed cropping with gramineous species on micronutrient concentrations and iron chlorosis of peanut plants grown in a calcareous soil. Plant Soil. 2008; 36:23–36.

12. Fang Q, Yu Q, Wang E, Chen Y, Zhang G, Wang J, et al. Soil nitrate accumulation, leaching and crop nitrogen use as influenced by fertilization and irrigation in an intensive wheat–maize double cropping system in the North China Plain. Plant Soil. 2006; 284: 335–350.

13. Li L, Li SM, Sun JH, Zhou LL, Bao XG, Zhang HG, et al. Diversity enhances agricultural productivity via rhizosphere phosphorus facilitation on phosphorus-deficient soils. Proc Natl Acad Sci. 2007; 104: 11192–11196.

14. Burns RG, DeForest JL, Marxsen j, Sinsabaugh RL, Stromberger ME, Wallenstein MD, et al. Soil enzymes in a changing environment:current knowledge and future directions. Soil Biol. Biochem. 2013; 58: 216–234.

15. Pignataro A, Moscatelli MC, Mocali S, Grego S, Benedetti A. Assessment of soil microbial functional diversity in a coppiced forest system. Appl Soil Ecol. 2012; 62: 115–123.

16. Qian X, Gu J, Pan HJ, Zhang KY, Sun W, Wang XJ, et al. Effects of living mulches on the soil nutrient contents, enzyme activities, and bacterial community diversities of apple orchard soils. Eur J Soil Biol. 2015; 70: 23–30.

17. Kourtev PS, Ehrenfeld JG, Haggblom M. Experi-mental analysis of the exotic and native plant species on the structure and function of soil microbial communities. 2003; 35: 895–905.

18. Baudoin E, Benizri E, Guckert A. Impact of artificial root exudates bacterial community structure in bulk soil and maize rhizosphere. Soil Biol Biochem. 2003; 35: 1183–1192.

19. Bainard LD, Koch AM, Gordon AM, Klironomos JN. Growth response of crops to soil microbial communities from convention monocrropping and tree-based intercropping systems. Plant Soil. 2013, 363:345–356.

20. Sun YM, Zhang NN, Wang ET, Yuan HL, Yang JS, Chen WY. Influence of intercropping and intercropping plus rhizobial inoculation on microbial activity and community composition in rhizosphere of alfalfa (*Medicago sativa* L.) and Siberian wild rye (*Elymus sibiricus* L.). FEMS Microbiol Ecol. 2009; 70 (2) :218–226.

21. Dai CC, Chen Y, Wang XX, Li PD. Effects of intercropping of peanut with the medicinal plant Atractylodes lancea on soil microecology and peanut yield in subtropical China. Agroforest syst. 2013; 87: 417–426.

22. Zuo Y, Zhang f. Iron and zinc biofortification strategies in dicot plants by intercropping with gramineous species. A review. Agron Sustain Dev. 2009; 29: 63–71.

23. Polthanee A, Wanapat S, Mangprom P. Row arrangement of peanut in cassava-peanut intercropping:II. Nutrient removal and nutrient balance in soil. Khon Kaen Agric J. 1998; 26 (3): 125–131

24. Kotchasatit A. Growth, yield and nutrient uptake of cassava and peanut in cassava/peanut intercropping systems under rained conditions at Khon Kaen Province. M.S Thesis, Faculty of Agriculture, Khon Kaen University. 1999.

25. Guan SY. Soil enzyme and its research method, China Agriculture Press, Beijing, China. 1986; pp. 274–340.

26. Yao HY, Huang CY. Soil microbial ecology and experimental technology, Science Press, Beijing, China. 2006; pp. 138–142.

27. Du S, Gao XZ. Technical specification for soil analysis, China Agriculture Press, Beijing, China. 2006; 15–163.

28. Bergmann GT, Bates ST, Eilers KG, Lauber CL, Caporaso JG, Walters WA, et al. The under-recognized dominance of Verrucomicrobia in soil bacterial communities. Soil Biol Biochem. 2011; 43(7):1450–1455.

29. Caporaso JG, Lauber CL, Walters WA, Berg-Lyons D, Lozupone CA, Turnbaugh PJ, et al. Global patterns of 16S rRNA diversity at a depth of millions of sequences per sample. Proc Natl Acad Sci USA. 2011; 108 (1): 4516–4522.

30. Magoč T, Salzberg SL. FLASH:fast length adjustment of short reads to improve genome assemblies. Bioinformatics. 2011; 27(21): 2957–2963.

31. Caporaso JG, Kuczynski J, Stombaugh J, Bittinger K, Bushman FD, Costello EK, et al. QIIME allows analysis of high-throughput community sequencing data. Nat Methods. 2010; 7(5): 335–336.

32. Wang Q, Garrity GM, Tiedje JM, Cole JR. Naive Bayesian classifier for rapid assignment of rRNA sequences into the new bacterial taxonomy. Appl Environ Microbiol. 2007; 73(16): 5261–5267.

33. Schloss PD, Westcott SL, Ryabin T, Hall JR, Hartmann M, Hollister EB, et al. Introducing mothur:open-source, platform-independent, community-supported software for describing and comparing microbial communities.Appl Environ Microbiol. 2009; 75(23): 7537–7541.

34. Li XP, Mu YH, Cheng Y, Liu XG, Nian H. Effects of intercropping sugarcane and soybean on growth, rhizosphere soil microbes, nitrogen and phosphorus availability. Acta Physiol Plant. 2013; 35(4): 1113–1119.

35. Ma YH, Fu SL, Zhang XP, Zhao K, and Chen H. Intercropping improves soil nutrient availability, soil enzyme activity and tea quantity and quality. Appl Soil Ecol. 2017; 119: 171–178.

36. Zhang JE, Gao AX, Xu HQ, Luo MZ. Effects of maize/peanut intercropping on rhizosphere soil microbes and nutrient content. Chin J Appl Ecol. 2009; 20(7): 1597–1602.

37. Li JL, Tu PF, Chen N, Tang JC, Wang XR, Nian H, et al. Effects of tea intercropping with soybean. Sci Agric Sin. 2008; 41(7): 2040–2047.

38. Xu Q, Cheng ZH, Meng HW, Zhang Y. Relationships between soil nutrients and rhizospheric soil microbial communities and enzyme activities in a maize-capsicum intercropping system. Chin J Appl Ecol. 2007; 18(12): 2747–2754.

39. Shi WM. Root exudates and soil nutrient availability. Soil. 1993; 25(5): 252–256.

40. Vander Heijden MGA, Wagg C. Soil microbial diversity and agro-ecosystem functioning. Plant Soil. 2013; 363: 1–5.

41. Zhang XQ, Huang GQ, Bian XM, Zhao QG. Effects of nitrogen fertilization and root interaction on the agronomic traits of intercropped maize, and the quantity of microorganisms and activity of enzymes in the rhizosphere. Plant Soil. 2013; 368(1-2): 407–417.

42. Kuang SZ, Tian SY, Li CY, Yi GJ, Peng Q. Effect of banana intercropping pattern and straw compost-returnon soil enzyme activity. Chin J Eco-Agric. 2010; 18(3): 617–621.

43. Chai Q, Huang P, and Huang GB. Effect of intercropping on soil microbial and enzyme activity in the rhizosphere. Acta Prataculturae Sinica. 2005; 14(5): 105–110.

44. Cai Q, Du GD, Lv DG, He Y, Jiang T, Sun JJ, et al. Effect of different kinds of intercropping with fruit/grass (crop) on soil microorganisms and enzymes in the south Horqinsan dyland. Agricultural Research in the Arid Areas. 28(4): 217–222.

45. Kennedy, A.C., and K.L. Smith. 1995. Soil microbial diversity and the sustainability of agricultural soils. Plant Soil. 2010; 170: 75–86.

46. Wu H, Kong Y, Yao YC, Bi NN, Qi LP, Fu ZG. Effects of intercropping aromatic plants on soil microbial quantity and soil nutrients in pear orchard. Sci Agric Sin. 2010; 43(1): 140–150.

47. Wang Y, Meng YL, Chen BL, Zhou ZG, HM Shu, HY Bian. Studies on the soil microorganism quality and soil nutrient content at the rhizosphere and non-rhizosphere region of cotton in wheat-cotton intercropping system. Sci Agric Sin. 2006; 26(10): 3485–3490.

48. Lu YH, Zhang FS. The advances in rhizosphere microbiology. Soil. 2006; 38 (2): 113–121.

49. Dong Y, Tang L, Zheng Y, Zhu YY, Zhang FS. Effects of nitrogen application rate on rhizosphere microbial community in wheat-faba bean intercropping system. Chin J Appl Ecol. 2008; 19(7): 1559–1566.

50. Song YN, Marschner P, Zhang FS, Bao XG, Li L. Effect of intercropping on bacterial community composition in rhizoshpere of wheat(*Triticum aestivum* L.), maize (*Zea mays* L.) and faba bean (*Viciafaba* L.). Acta Ecol Sin. 2006; 26(7): 2268–2274.

51. Ren TZ, Grebo S. Soil bioindicators in sustainable agriculture. Sci Agric Sin. 2000; 33(1): 68–75.

52. Ma DY, Guo TC, Song X, Wang CY, Zhu YJ, Wang YH, et al. Effects of urea application rate on the quantity of microorganisms and activity of enzymes in wheat rhizosphere. Acta Ecol Sin. 2007; 12: 5222–5228.

53. Fierer N, Leff JW, Adams BJ, Nielsen UN, Bates ST, Lauber CL, et al. Cross-biome metagenomic analyses of soil microbial communities and their functional attributes. Proc Natl Acad Sci USA. 2012; 109: 21390–21395.

54. Li X, Sun M, Zhang H, Xu N, Sun G. Use of mulberry–soybean intercropping in salt–alkali soil impacts the diversity of the soil bacterial community. Microb Biotechnol. 2016; 9(3):293.

55. Yang S, Wen X, Jin H, Wu Q. Pyrosequencing investigation into the bacterial community in permafrost soils along the China-Russia crude oil pipeline (CRCOP). PLoS One. 2012; 7: e52730.

56. Kersters K, Vos P De, Gillis M, Swings J, Vandamme P, Stackebrandt E. Introduction to the Proteobacteria. In:Martin D (ed) The prokaryotes, vol 5, Proteobacteria: Alpha and Beta SubclassesSpringer, New York. 2006; 3–37.

57. Shang SH, Yi YL. A greenhouse assay on the effect of applied urea amount on the rhizospheric soil bacterial communities. Indian J Microbiol. 2015, 55(4): 406–414.

58. Brons JK, Van Elsas JD. Analysis of bacterial communities in soil by use of denaturing gradient gel electrophoresis and clone libraries, as influenced by different reverse primers. Appl Environ Microbiol. 2008; 9: 2717–2727.

59. Kielak A, Rodrigues JLM, Kuramae EE, Chain PSG, Van Veen JA, Kowalchuk GA. Phylogenetic and metagenomic analysis of Verrucomicrobia in former agricultural grassland soil. FEMS Microbiol Eco. 2010; 171: 23–33.

60. Wang G, Jin J, Liu J, Chen X, Liu J, Liu X. Bacterial community structure in a mollisol under long-term natural restoration, cropping and bare fallow history estimated by PCR-DGGE. Pedosphere. 2009; 19: 156–165.

61. Shen C, Ge Y, Yang T, Chu H. Verrucomicrobial elevational distribution was strongly influenced by soil ph and carbon/nitrogen ratio. J Soil Sediment. 2017; 17(10): 1–8.

62. Zhang H, Sekiguchi Y, Hanada S, Hugenholtz P, Kim H, Kamagata Y, et al. *Gemmatimonas aurantiaca* gen nov, sp nov, a Gram-negative, aerobic, polyphosphate-accumulating micro-organism, the first cultured representative of the new bacterial phylum Gemmatimonadetes phyl. nov. Int J Syst Evol Microbiol. 2003; 53: 1155–1163.

63. Hanada S, Sekiguchi Y. The Phylum Gemmatimonadetes. In: Rosenberg E, DeLong E F, Lory, S., E. Stackebrandt, F. Thompson (eds) The Prokaryotes. Springer, Berlin, Heidelberg. 2014.

64. He Z, M Xu, Y Deng, S Kang, L Kellogg, L Wu, et al. Metagenomic analysis reveals a marked divergence in the structure of belowground microbial communities at elevated CO_2_. Ecol Lett. 2010; 13: 564–575.

65. He Z, Piceno Y, Deng Y, Xu M, Lu Z, DeSantis T, et al. The phylogenetic composition and structure of soilmicrobial communities shifts in response to elevated carbon dioxide, ISME J. 2012; 6: 259–272.

66. Xu M, Chen X, Qiu M, Zeng X, Xu J, Deng D, et al. Bar-coded pyrosequencing reveals the responses of PBDE-degrading microbial communities to electron donor amendments. PLoS One. 2012; 7: e30439.

67. Zhang W, Chen L, Zhang R, Lin K. High throughput sequencing analysis of the joint effects of bde209-pb on soil bacterial community structure. J Hazard Mater. 2016; 301: 1–7.

68. Brewer TE, Handley KM, Carini P, Gilbert JA, Fierer N. Genome reduction in an abundant and ubiquitous soil bacterium ‘candidatus udaeobacter copiosus’. Nat Microbiol. 2016; 2: 16198.

